# Properties of decision-making tasks govern the tradeoff between model-based and model-free learning

**DOI:** 10.1101/730663

**Authors:** Zachary M Abzug, Marc A Sommer, Jeffrey M Beck

## Abstract

When decisions must be made between uncertain options, optimal behavior depends on accurate estimations of the likelihoods of different outcomes. The contextual factors that govern whether these estimations depend on model-free learning (tracking outcomes) vs. model-based learning (learning generative stimulus distributions) are poorly understood. We studied model-free and model-based learning using serial decision-making tasks in which subjects selected a rule and then used it to flexibly act on visual stimuli. A factorial approach defined a family of behavioral models that could integrate model-free and model-based strategies to predict rule selection trial-by-trial. Bayesian model selection demonstrated that the subjects strategies varied depending on lower-level task characteristics such as the identities of the rule options. In certain conditions, subjects integrated learned stimulus distributions and tracked reward rates to guide their behavior. The results thus identify tradeoffs between model-based and model-free decision strategies, and in some cases parallel utilization, depending on task context.

## 1. Introduction

Humans are often forced to make decisions between uncertain options, where the probability of each option leading to success is unknown and must be estimated in order to behave optimally. For example, imagine you are given the choice between shooting a basketball into a net or a soccer ball into a goal, but the distance from which you would have to take each shot is unknown (Fig. 1A). Model-*free* reinforcement learning (RL) algorithms would solve this problem by tracking past outcomes (successes and failures) for each possible action (soccer and basketball) to determine the action with the greater expected probability of success. However, sensory cues can also carry important information about expected success probabilities: basketball shots from halfcourt should be harder than shots from the three-point line, and both should be harder than shooting a soccer ball from only six yards away. Model-*based* RL algorithms utilize these sensory cues to estimate the generative processes that control the probability of success for any given action. In our example, a model-based decision-maker would utilize sensory data from previous trials (Fig. 1B) to learn the probability distributions over possible shooting distances for both soccer and basketball (Fig. 1C).

**Figure 1:**
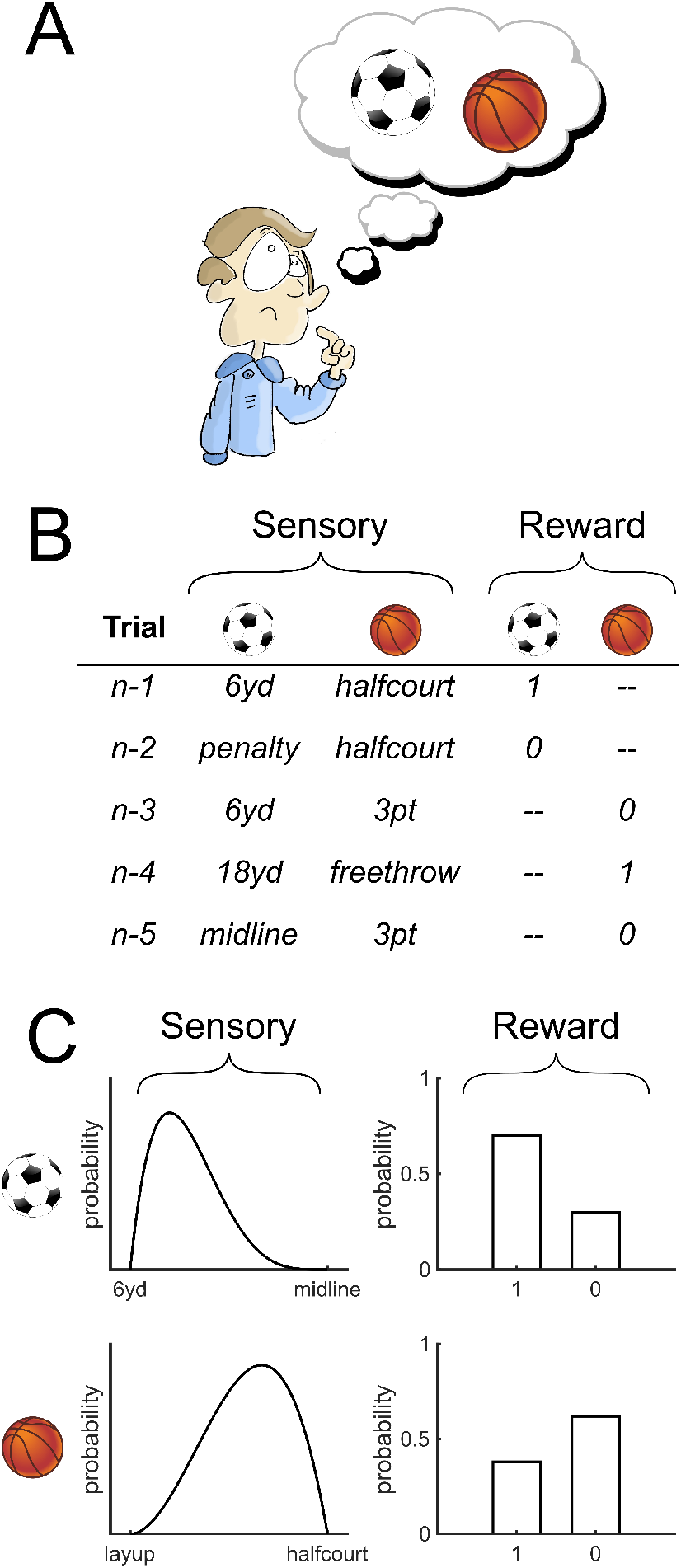
Multiple sources of information affect decision-making under uncertainty. (A) Often, subjects have to make decisions between two uncertain tasks, for instance, whether to shoot a basketball or a soccer ball from an unknown distance. (B) To decide which option is most likely to result in reward, subjects can utilize both sensory information (the distances that past shots have been taken from) and reward information (the outcomes of past shots). (C) These two sources of information can be used to estimate the generative process underlying each task (the distributions of shot distances, left) or the action-values associated with each task (the probability of reward, right).

Many lines of research have demonstrated that, in certain contexts, model-based RL algorithms that incorporate estimates of generative processes outperform model-free algorithms that rely solely on tracking outcomes [1–3]. However, neurophysiological studies of decision-making have tended to focus on correlates of model-free variables (such as reward prediction error) rather than model-based variables [4–6]. Recently, researchers have found that many of the same brain regions putatively involved in model-free RL, such as the dopaminergic system [7], also seem to underlie model-based RL (see 8 for review).

Implicit in model-based RL is the statistical learning of probability distributions over task relevant stimuli. Humans are capable of implicitly learning Gaussian, skewed unimodal, and bimodal distributions [9–14], although the learning of skewed or multimodal distributions takes longer and is less accurate overall [9]. Crucially, humans are also able to update their beliefs and learn non-static distributions to make decisions in changing environments [15–19] and even identify correlations between the outcomes of different choices [1].

Substantial research has also studied how humans choose between different options after the probabilities of reward associated with each option have already been estimated. Studies often report that humans perform “probability matching” by choosing different options in proportion to each option’s relative probability of leading to reward [20–22]. However, alternate lines of work have demonstrated that probability matching arises due to a mismatch between the generative processes that actually determine reward probabilities and subjects’ beliefs about those generative processes [1]. Furthermore, in certain cases, such as those where reward probabilities are partially dependent on the subject’s motor behavior, subjects exhibit behavior consistent with the maximization of reward rates rather than probability matching [1, 9].

In this study, our goal was to determine how subjects utilize different sources of information to make decisions under uncertainty, and how this utilization depends on the specific decisions required. To this end, we developed two serial decision-making tasks that required subjects to first pre-select a behavioral rule for a subtask in which they applied that rule to flexibly act on visual stimuli. We also developed a family of behavioral models that estimated the difficulty of each rule and predicted subjects’ trial-by-trial rule selections as a function of two sources of information: the observed patterns of past visual stimuli and rewards. In a prior study, we demonstrated that on a session-by-session basis, monkeys’ rule preferences in a similar serial decision-making task were correlated with their relative performance on each rule [23]. In the present study, we used human subjects to extend this work and found that when task difficulty systematically changes over the course of a behavioral session, the subjects used both sensory and reward information to estimate task difficulty. The degree to which sensory vs. reward information was utilized depended on the specific decisions required, with underlying strategies, accordingly, trading off between model-based and model-free behaviors, and sometimes implementing them in parallel.

## 2. Results

In this study, we developed two serial decision-making tasks that shared a common structure but required humans to make different specific decisions: subjects first pre-selected a behavioral rule, then used that rule to flexibly act on visual stimuli. We then used factorial modeling [9, 24] to generate a large family of candidate behavioral models and compared their performance across groups of subjects. Models were fit to subjects’ rule selections on a trial-by-trial basis based upon their past history of observed sensory (i.e., visual) features as well as trial outcomes. Model performance was assessed via Bayesian Model Selection [25] and rule selection prediction to identify decision-making strategies.

In both behavioral tasks, subjects were first shown two or three colored stimuli, each corresponding to an abstract behavioral rule according to a fixed color-rule contingency on which subjects had been previously instructed. On each trial, subjects could freely select which rule they wanted to implement in the upcoming second stage of the trial. After rule selection was done, visual stimuli appeared that varied in rule-relevant ways. For instance, in Experiment 1 (Fig. 2), where the behavioral rules were “pick the smaller target” and “pick the darker target”, visual stimuli could vary in both their sizes and their brightnesses. Subjects then implemented their chosen rule, and received feedback indicating whether they had implemented the rule correctly or incorrectly. On each trial, the subjects were presented with two sources of information that could be used to guide their future rule selections: the sensory features of the visual stimuli that indicated how difficult each rule was on that trial, and feedback that indicated whether the selected behavioral rule was performed correctly. Both of these sources of information were of potential importance, because the objective difficulties of each rule (i.e., the distributions from which the size and brightness differences were drawn from) dynamically varied across the behavioral session.

**Figure 2:**
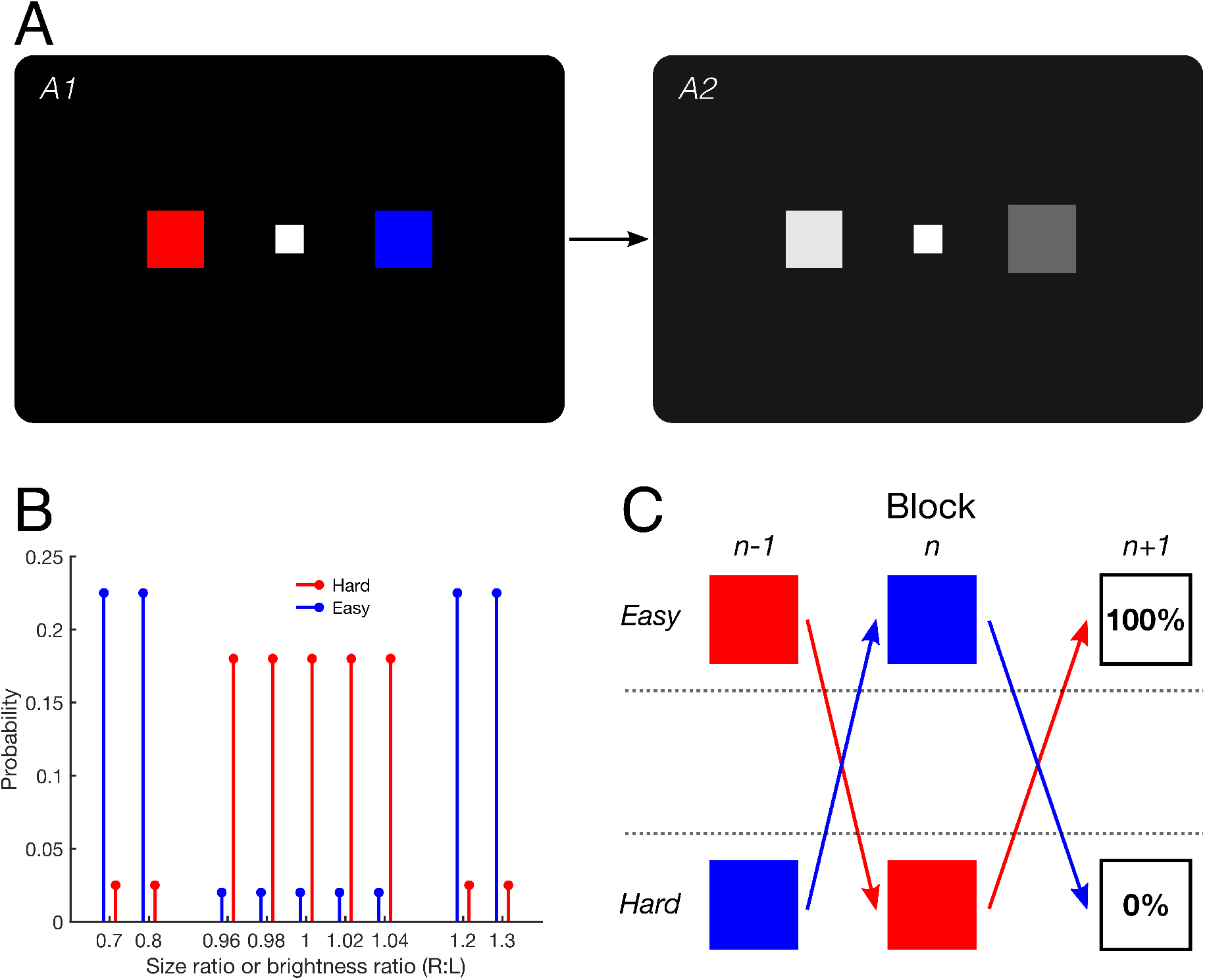
Task schematic for Experiment 1. (A) Subjects used button presses to select one of two colored rule targets (*A1*). Subjects were previously instructed of the relationship between target color and behavioral rule. For example, the red target could correspond to the size rule *R_S_* and the blue rule could correspond to the brightness rule *R_B_*. They were then shown two visual stimuli, differing in their sizes and brightnesses (*A2*). The subject’s goal was, using a button press, to select the correct target given the previously selected rule. Stimuli not to scale; they are shown larger than in the experiment for clarity of illustration. (B) On each trial, each rule was classified as “easy” or “hard”. Actual size and brightness ratios were drawn from overlapping discrete distributions. Each difficulty distribution was a mixture of discrete uniform distributions corresponding to ratios near unity (*n* = 5 values, ***π*** = 0.1 for “easy” distribution, ***π*** = 0.9 for “hard” distribution) and ratios significantly above or below unity (*n* = 4 values, ***π*** = 0.9 for “easy” distribution, ***π*** = 0.1 for “hard” distribution). Red and blue bars are visually offset to avoid overlap. See Methods for further details. (C) Trials were organized into a block structure. At the end of each block, the rules swapped difficulty classes with a 100% contingency. For instance, if the rule denoted by the red target was hard in block *n*, it would be easy in block *n* + 1 (red arrow).

### 2.1. Behavioral models of serial decision-making tasks

Overall, our objective was to model subjects’ rule selection strategies across trials in each of our two experiments. We hypothesized that subjects would track both sensory features (e.g., the size and brightness differences between decision stimuli in Experiment 1) and feedback across past trials, and integrate those two sources of information in order to guide their rule selection on subsequent trials. Therefore, our core behavioral model featured three modules: the Sensory Module, the Reward Module, and the Decision Module (Fig. 3). We then used a factorial modeling approach to perturb our core model and generate a large family of candidate behavioral models to predict rule selections. We varied the following four model factors:

1. *Decision Model* (2 levels): stochastic posterior model (‘SPK’) or posterior probability matching (‘PPM’)
2. *Sources of Information* (4 levels): both sensory and reward information (‘SR’), sensory information only (‘S’), reward information only (‘R’), or both sources with matched learning rates (‘SRM’)
3. *Decision Bias* (2 levels): present (‘B’) or absent (‘NB’)
4. *Lapse* (2 levels): present (‘L’) or absent (‘NL’)

**Figure 3:**
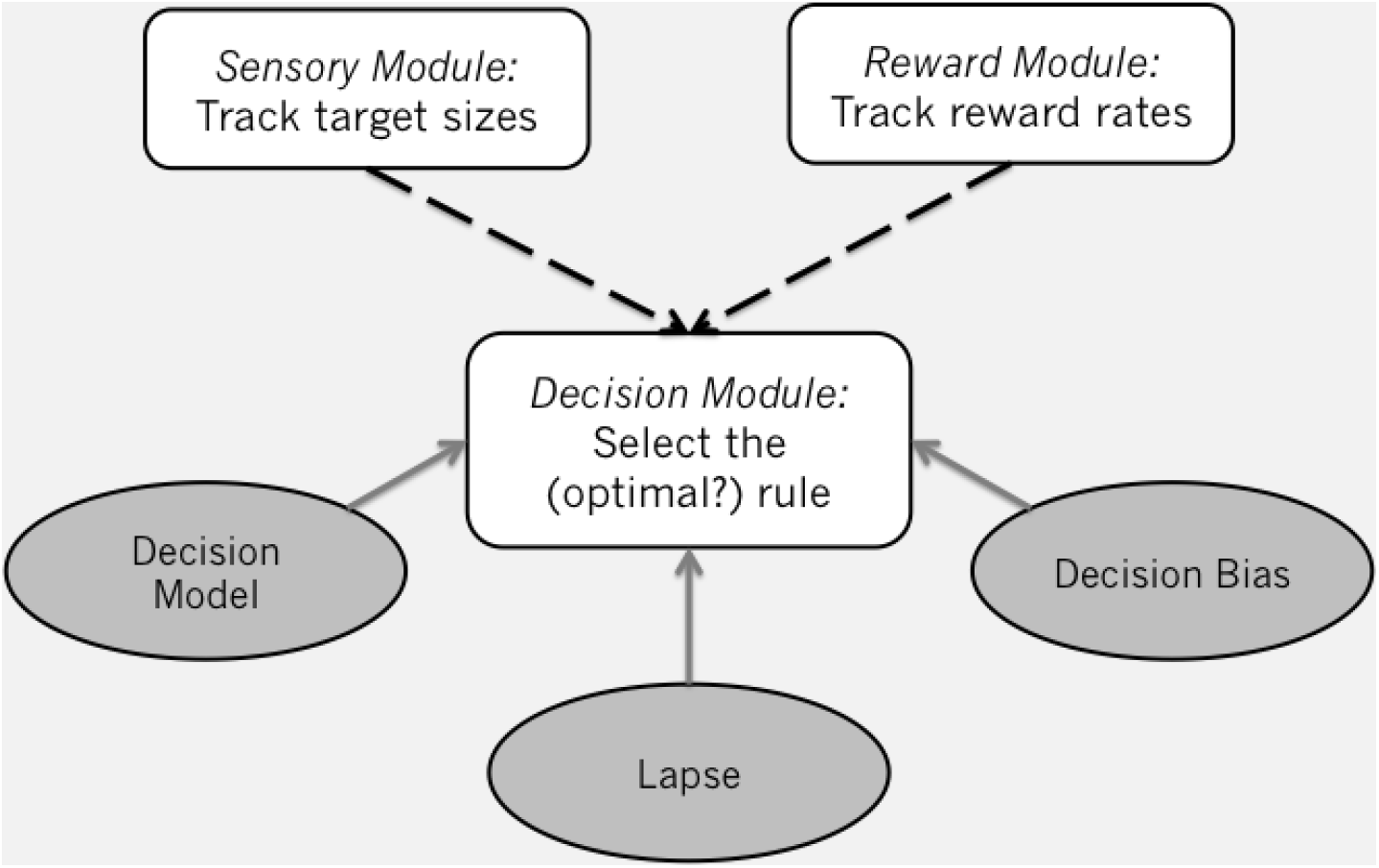
Generating the family of models. We first constructed a model that tracked both sensory features and feedback across past trials, and integrated those two sources of information in order to guide rule selection on subsequent trials. We then perturbed the model systematically to generate a family of 32 models by changing four factors: the optimality of the *Decision Model*, whether both *Sensory* and *Reward* information were used (dashed lines), and whether the subjects’ *Decision Bias* and *Lapse* rate were incorporated into the model.

By selecting levels for each of these four factors we could generate a behavioral model and a unique string to describe it. For example, a model that implements probability matching, uses only sensory information, and has decision bias but no lapse can be denoted as ‘PPM-S-B-NL’. By taking every possible combination of levels, we generated 32 candidate behavioral models, each with their own identifying string. We also included two control models, ‘BiasOnly’ and ‘SensoryChaser’, yielding 34 models in total. The ‘BiasOnly’ model does not use either sensory or reward information, and uses a static decision model on each trial that represents the subjects’ learned biases. The ‘SensoryChaser’ model simulates the behavior of a subject who always selects the rule that was easiest on the previous trial with some learned probability *p*. The factors and their levels are discussed briefly below. For more details on all models, see Methods.

#### Decision Model

Many studies of probabilistic decision-making have reported that subjects sample different options in proportion to their relative probabilities of reward [20–22]. This behavior, referred to as “probability matching” or “posterior probability matching” (‘PPM’), is equivalent to sampling from the posterior distribution and is suboptimal: the optimal strategy is to always select the option with the highest probability of reward. The power function approximation to the stochastic posterior model [stochastic posterior, *κ*-power; ‘SPK’; 9] increases flexibility by raising the posterior distribution to some power *κ* prior to sampling. With *κ* = 1, the ‘PPM’ and ‘SPK’ models are equivalent. However, with *κ* > 1, the ‘SPK’ model approaches optimality by increasing the probability of selecting options with high reward probabilities. In the limit *κ* → ∞, the model simply implements the decision believed to maximize reward..

#### Sources of Information

We explicitly varied whether our behavioral models tracked sensory features, reward history, or both. The most flexible model tracks both sensory and reward information and uses both sources of information to guide rule selections (‘SR’). Suboptimalities can be induced by matching the learning rates across the Sensory and Reward Modules (‘SRM’), only utilizing sensory information (‘S’), or only utilizing reward information (‘R’).

#### Decision Bias

Suboptimalities also could be introduced into the models by explicitly modeling decision biases (‘B’). These biases represent innate preferences for particular behavioral rules that are not explained by the history of observed sensory and reward information. Models with no biases (‘NB’) have less behavioral flexibility.

#### Lapse

Lastly, suboptimalities can be introduced by including behavioral lapses (‘L’). Lapses are often included in behavioral models to explain decisions where subjects select a suboptimal option even though the decision is objectively easy, perhaps because of a lapse of attention on some subset of trials. Models with no lapses (‘NL’) have less behavioral flexibility.

#### 2.1.1. Comparing model fits across the population

In each experiment, we fit data from each behavioral session with all 34 candidate models, and calculated the Bayesian information criterion (*BIC*) for each model. *BIC* is a measure of model performance that accounts for the complexity of the model and can be used to compare model performance *within* a behavioral session. We also used Bayesian model selection [BMS; 25] to determine which models and factor levels best explained rule selections across the population. The particular BMS algorithm we used is a variational Bayesian algorithm that uses marginal likelihoods (or an approximation to the marginal likelihood, such as (−1/2)*BIC*) to compare model performance *across* behavioral sessions or subjects, by accounting for in-group heterogeneity. Lastly, we looked at predictions accuracy *across* models: the proportion of trials that the model selects the same rule that the subject selected. In most cases, we looked at prediction accuracy relative to chance, which was defined as prediction accuracy for the ‘BiasOnly’ model.

### 2.2. Experiment 1

Twenty-four naïve human subjects completed an average of 465 ± 70.5 trials in an hourlong behavioral session. Each session consisted of blocks of 30-60 trials.

The task used in Experiment 1 is shown in Figure 2A. In general, subjects first had to select between two behavioral rules, each denoted by a colored stimulus (Fig. 2A, left). The two possible behavioral rules were “pick the smaller target” (size rule *R_S_*) and “pick the darker target” (brightness rule *R_B_*). On each trial, both rule options were presented, and their locations were counterbalanced across trials. After a rule stimulus was selected, subjects had to use that behavioral rule to flexibly discriminate between squares differing in their sizes and brightnesses (Fig. 2A, right). On each trial, the target sizes and brightnesses were probabilistically drawn from pre-determined distributions (Fig. 2B). If the subject correctly performed the perceptual discrimination in accordance with the previously selected (or instructed) rule, they received positive feedback.

Transitions between blocks were continuous, covert, and uncued. Within a single block, rule difficulties were globally coupled: one rule was designated as the “easy” rule and the other rule was necessarily designated as the “hard” rule (these assignments swapped at the start of every new block; Fig. 2C). Importantly, the difficulty distributions were overlapping, such that a single trial could, for example, feature both a difficult size discrimination and a difficult brightness discrimination. For more details on how target sizes and brightnesses were generated on a trial-by-trial basis, see Methods.

#### 2.2.1. Results & Discussion

We fit our set of 34 behavioral models to each of 24 human subjects and used the *BIC* and BMS to determine the best-fitting model for each subject while taking into account model complexity. Figure 4 shows the Δ*BIC* for each model fit, relative to the best-fitting model for each subject, where smaller Δ*BIC* values (warmer colors) correspond to better fits. The models are ordered according to the posterior probability that the decision model was utilized by a randomly chosen subject, as determined by BMS. Most subjects were well-fit by multiple models, without any models clearly outperforming the rest at the group level. Importantly, all models were at least as likely as our two control models, ‘BiasOnly’ and ‘SensoryChaser’. Five models had a > 0.05 probability of being the best-fitting model for a random subject, and no models had a > 0.2 probability of being the best-fitting model (Fig. 5A). These results are confirmed by Figure 5B: out of the four possible factors, only the lapse factor showed population-level effects, with random subjects being more likely to not have lapses (exceedance probability *P*\* > 0.999, see Methods for further details). Additionally, only 6.2% of subjects were likely to be described by an observer model using only sensory information (*P*\* = 0.0003).

**Figure 4:**
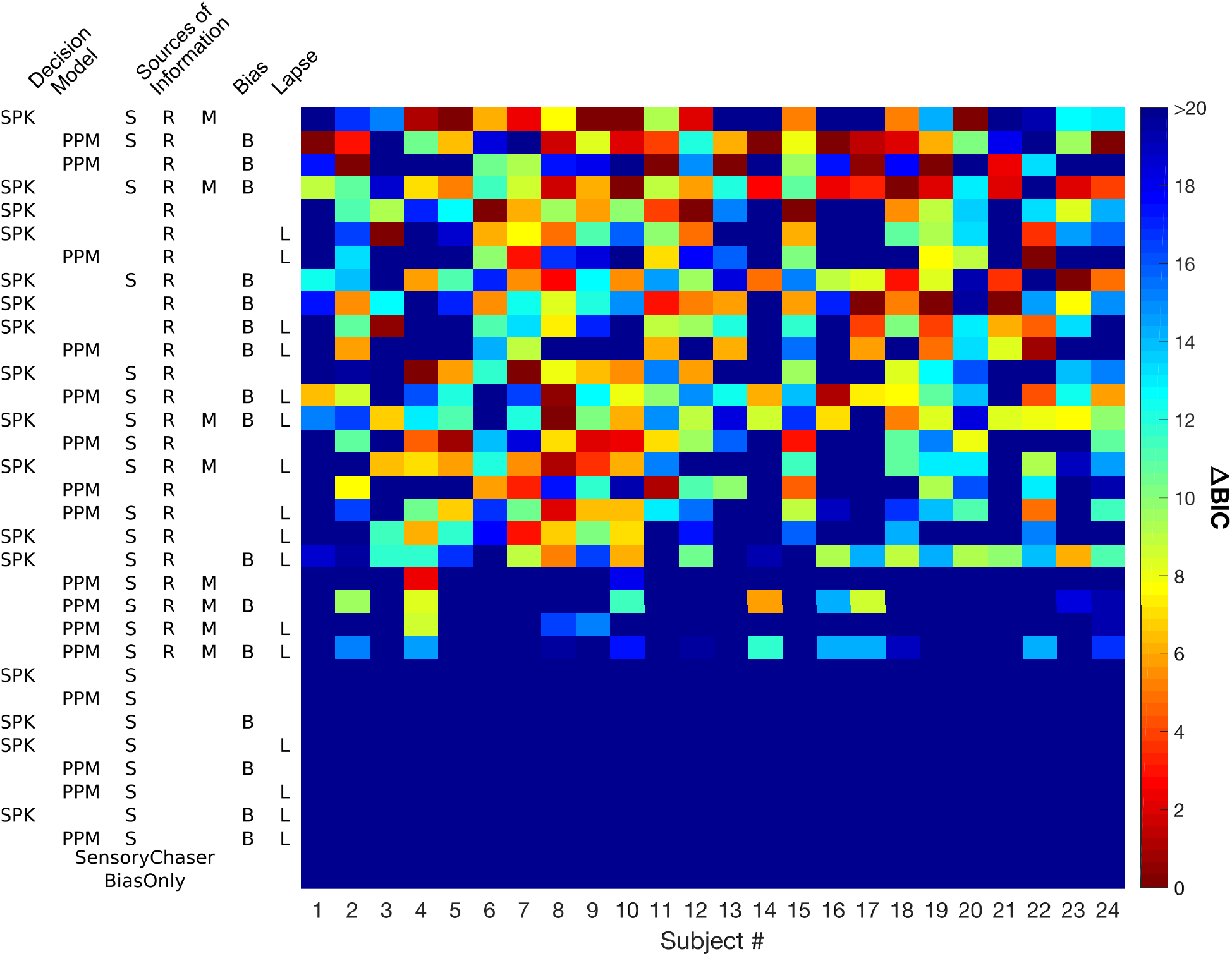
Model fitting results for Experiment 1. Each column represents a subject, and each row represents an observer model from the factorial family of models and two control models. Models are identified by a model string (substrings ‘NB’ and ‘NL’ are left out for clarity), and are ordered according to model posterior likelihood from BMS algorithm. Cell color represents model evidence, defined here as the difference between the *BIC* for that model and the *BIC* for the the best model for that subject. Differences of greater than 6 are considered to be strong evidence in favor of the model with lower *BIC*.

**Figure 5:**
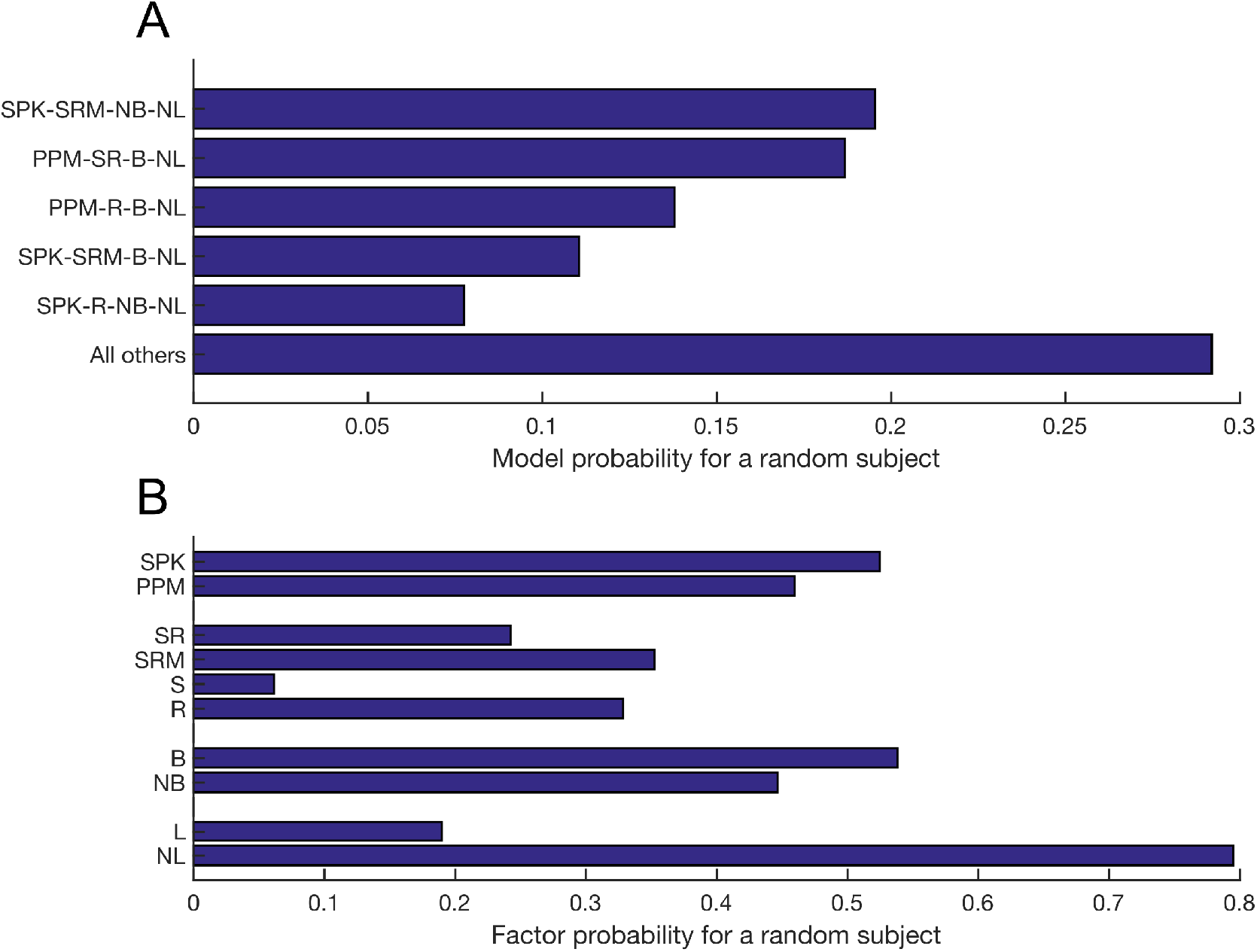
Model selection results for Experiment 1. (A) Probability that a given model generated the data of a randomly chosen subject. Only models with posterior likelihoods > 0.05 are shown individually. No individual models had posterior likelihoods > 0.2, and the top models have no obvious strong relation to each other. (B) Probability that a given model level within a factor generated the data of a randomly chosen subject. There was significantly more support for models without lapses than models with lapses (‘NL’ vs. ‘L’; *P*\* > 0.999). Observer models using only sensory information were unsupported (‘S’ vs. others; *P*\* = 0.0003).

In Experiment 1, 29/32 behavioral models predicted rule selections significantly better than the ‘BiasOnly’ model (Fig. 6; one-tailed t-test, Bonferroni-Holm correction for multiple comparisons), suggesting that subjects were using incoming sensory and/or reward information to guide their decisions. Unsurprisingly, the model with maximal complexity, ‘SPK-SR-B-L’, was the best at predicting rule selections across subjects (21.61 ± 10.2% above chance). However, this model was only one of 22 models with mean prediction improvements of 20% or greater, including all five of the top models as determined by BMS. Mean prediction improvements for each model were correlated with the number of parameters in the model (*r* = 0.515, *p* < 0.005) but not the BMS posterior probability for the model (*r* = 0.272, *p* = 0.12). As expected from the BMS results, models with lapses tended to predict rule selections better than models without lapses (paired *t*(15) = 1. 91, *p* = 0. 075), and models using only sensory information did not predict as well as models using reward information only or sensory and reward information together (one-way ANOVA on information source, *F*(3, 28) = 77.1; post-hoc Tukey’s test on ‘S’ vs. others, all *p* < 0.0001). Additionally, two other trends emerged: SPK models tended to predict better than PPM models (paired *t*(15) = 2.10, *p* = 0.053), and models with decision bias predicted better than those without (paired *t*(15) = 4.20, *p* < 0.001). In every case, as expected, more complex models were better at predicting behavior than less complex models.

**Figure 6:**
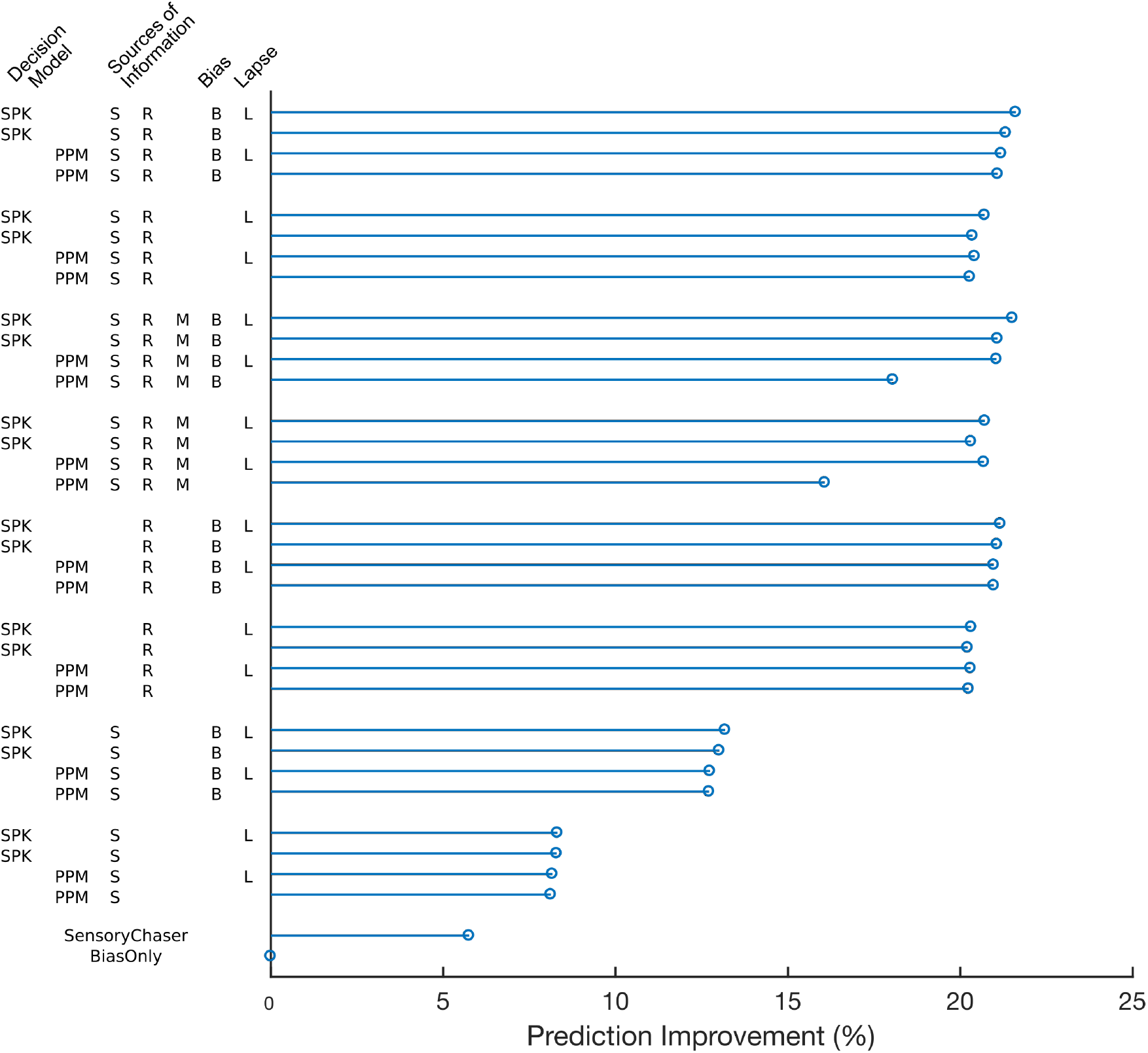
Prediction improvement for Experiment 1. Average prediction improvement was significantly above chance for all but 3/32 behavioral models, and was correlated with model complexity (*r* = 0.515, *p* < 0.005) but not BMS posterior probability (*r* = 0.272, *p* = 0.12).

Overall, the results of Experiment 1 suggest that subjects were using predominantly reward information to guide their rule selections, although the precise strategy used by each subject varied. All 32 models tested (and the ‘SensoryChaser’ control model) outperformed the ‘BiasOnly’ control model, as measured by both *BIC* and prediction improvement metrics, but no individual model or small subset of models significantly outperformed the rest of the family. Interestingly, unlike as is often found [e.g., 9, 26], models without lapses significantly outperformed models with lapses. One possible explanation for this result is that the incorporation of both decision biases and lapses are partially redundant when *N_R_* = 2, particularly when the lapse rate *ℓ* is small. Another complementary possible explanation has to do with the primary benefit of including lapse parameters: the ability to explain and compensate for errors on objectively easy (“high signal intensity”, e.g., high coherence levels on the RDK task) trials where proportion correct should approach or equal 1 [cf. 26, Fig. 1, solid light line vs. broken light line]. If signal intensities are not high enough for performance to saturate (in this study, for subjects to consistently select the optimal rule), then lapse parameters become much less important for accurate behavioral modeling.

### 2.3. Experiment 2

In Experiment 1, we found that the behavioral strategies used by individual subjects varied substantially across subjects. In Experiment 2, we sought to determine if there were certain task conditions in which behavioral strategies across subjects would converge. Additionally, we asked if we could promote the use of sensory information to guide decisionmaking by facilitating the comparison of sensory features across rules.

Specifically, we changed two key features of the task. First, all of our task rules operated on the same stimulus feature (target size) instead of distinct stimulus features. This means that subjects did not need to maintain and update multiple estimates of the relationships between a stimulus feature and reward probability, which should facilitate the comparison of sensory features across rules. Second, subjects implemented rules using fine motor behaviors instead of button presses, which should promote the maximization of reward rates over suboptimal probability matching strategies. This is thought to be due to subjects’ tendencies to attribute errors to motor errors rather than stimulus mis-estimation, thereby enhancing their confidence regarding the relationship between stimulus features and task difficulty [1]. Overall, we hypothesized that these changes would cause observed behavioral strategies across subjects to converge on strategies that tracked sensory information in near-optimal ways.

Twenty-four naïve human subjects (different from those used in Experiment 1) completed an average of 952 ± 164.2 trials in an hour-long behavioral session. In this experiment, we utilized a “shooting” variant of our serial decision-making task that required them to use a joystick to pre-select a color, and then shoot at a target of that color among multiple distractors (Fig. 7). A video of the task can be found at https://goo.gl/iXZybV. As in Experiment 1, behavioral sessions consisted of blocks, and transitions between blocks were continuous, covert, and uncued. Within a block, one colored rule was designated as the “easy” rule (i.e., largest target size, on average), one was designated as the “medium” rule, and the third was designated as the “hard” rule (i.e., smallest target size, on average; Fig. 7B). Blocks lasted 6-9 trials, and at the end of each block, there was a 40% chance of swapping the identities of the “easy” and “medium” rules, a 40% chance of swapping the identities of the “medium” and “hard” rules, and a 20% chance of the rankings staying the same (Fig. 7C). A single rule could never transition directly from “easy” to “hard” or vice versa in a single block transition. For more details on the task used in Experiment 2 and the motor behavior of subjects, see Methods.

**Figure 7:**
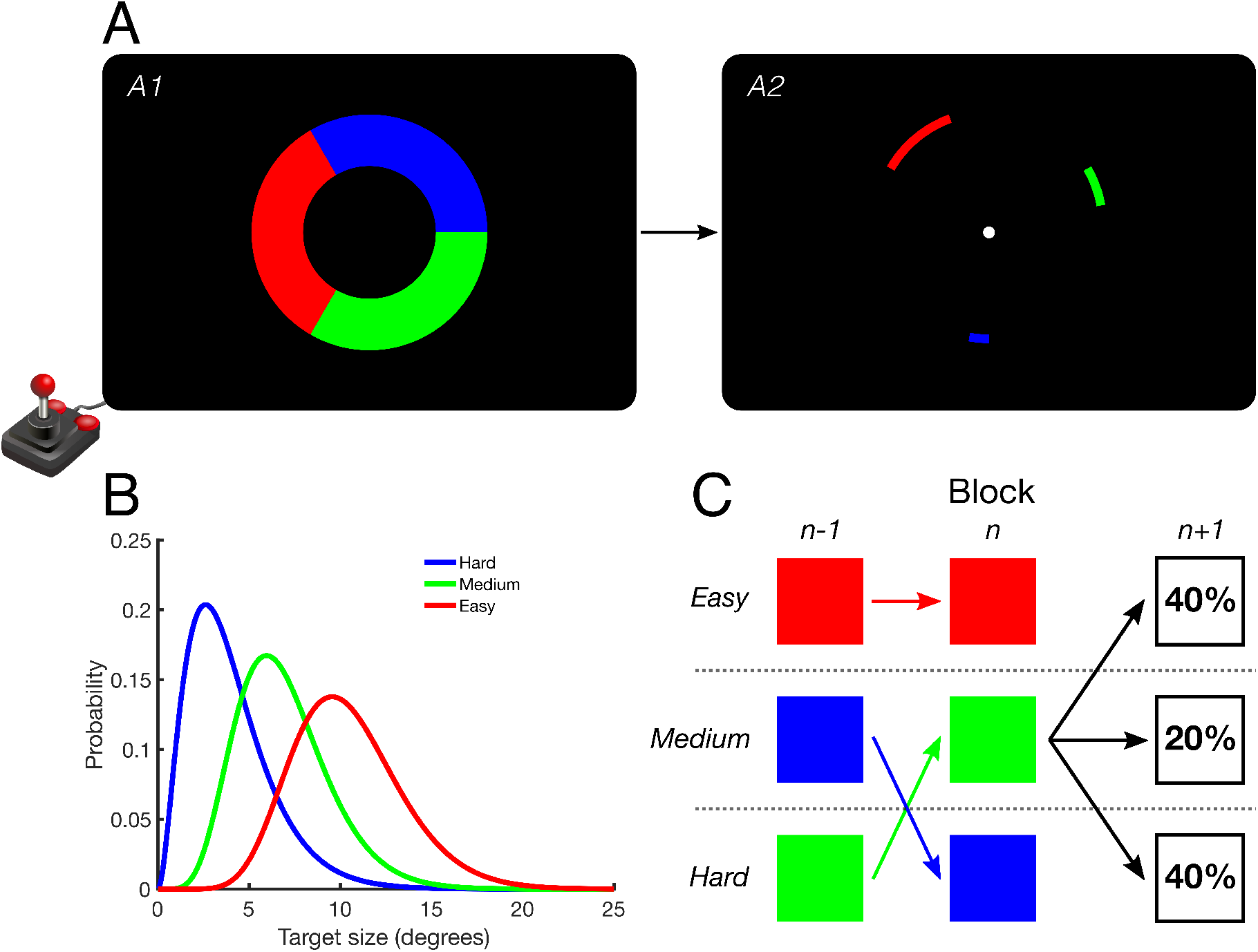
Task schematic for Experiment 2. (A) Subjects used a joystick to select one of three color rules (*A1*). They then used the joystick to shoot a virtual ball at the appropriately colored target, given the previously selected color rule (*A2*). A video of the task can be found at https://goo.gl/iXZybV. (B) On each trial, each color was classified as “easy”, “medium”, or “hard”. Actual target sizes were drawn from overlapping gamma distributions with class-specific parameters. (C) Trials were organized into a block structure. At the end of each block, the colored rules could swap difficulty classes. The “easy” and “medium” rules could swap (40%), the “medium” and “hard” rules could swap (40%), or the classes could stay the same (20%). The “easy” and “hard” rules could never directly swap in a single block transition.

#### 2.3.1. Results & Discussion

We fit our set of 34 behavioral models to each of 24 human subjects and used the *BIC* and BMS to determine the best-fitting model for each subject while taking into account model complexity. Figure 8 shows the Δ*BIC* for each model fit, relative to the best-fitting model for each subject. The models are ordered according to the posterior probability of each observer model from the BMS procedure. In contrast to Experiment 1, where most subjects were fit well by many different behavioral models, the model fits for most subjects in Experiment 2 were much more sparse and selective. Additionally, whereas few detectable patterns existed in the model rankings for Experiment 1, some clear patterns emerged in Experiment 2: the top two models (‘SPK-SR-NB-L’ and ‘SPK-SR-B-L’) are identical in three of the four factors of interest and only differ in the presence or absence of decision bias, and the third-best model (‘PPM-S-NB-NL’) is one of the simplest possible models in the family, with only one free parameter (*α_s_*). Additionally, in contrast to Experiment 1, where almost every subject was well-fit by a large number of models (the rectangular “block” of color in Fig. 4), subjects in Experiment 2 tended to either be fit well by many models (e.g., Subject 9) or very few models (e.g., Subject 13; vertical “stripes” of color in Fig. 8).

**Figure 8:**
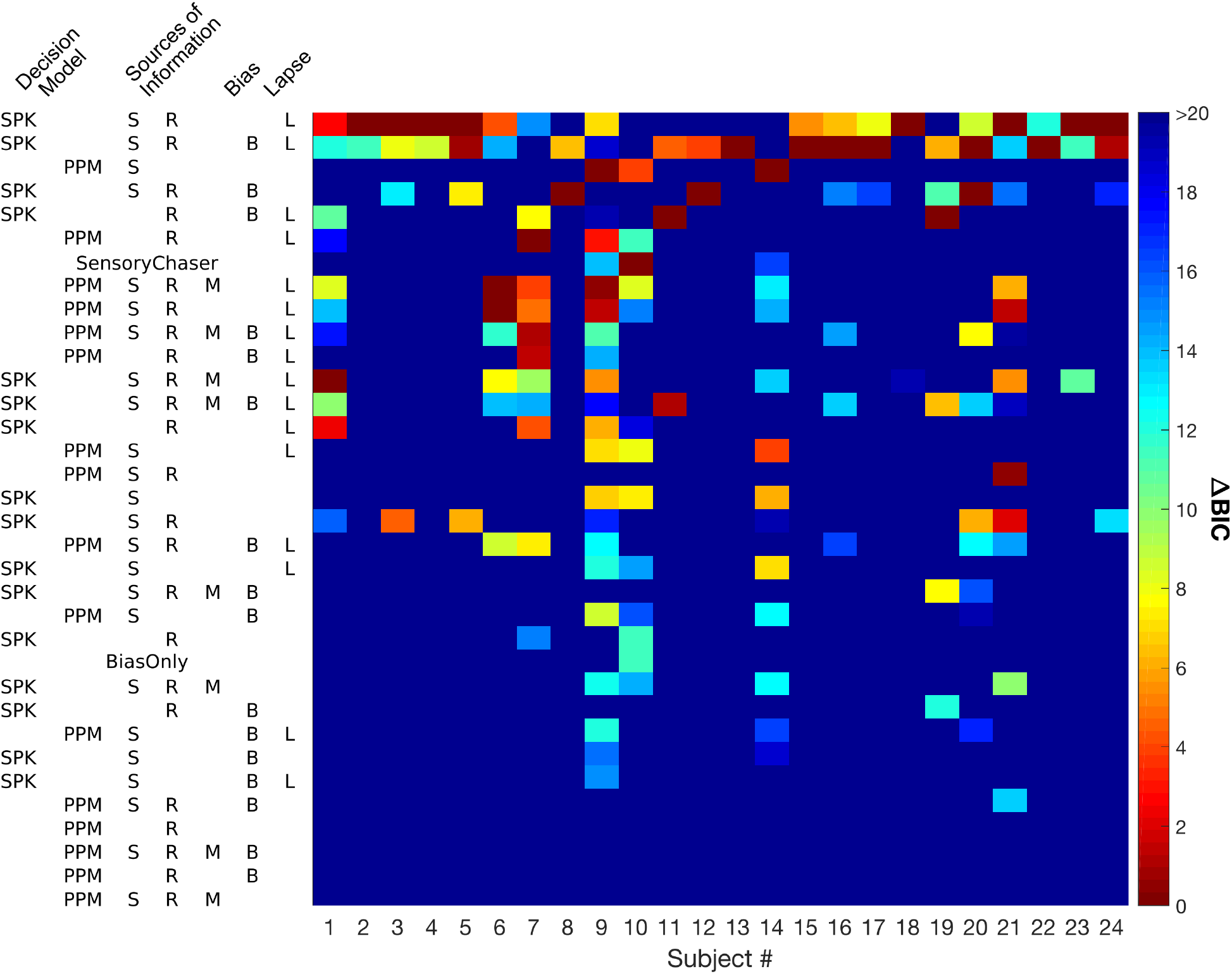
Model fitting results for Experiment 2. Same conventions as Fig. 4.

These patterns inferred from the Δ*BIC* values for each subject were corroborated by BMS over the population as a whole (Fig. 9). Observer models ‘SPK-SR-NB-L’ and ‘SPK-SR-B-L’ were the most likely to be used by a randomly selected subject, although neither model individually gained significance (Fig. 9A). Additionally, three of the four factors showed population-level effects (Fig. 9B). Subjects were significantly more likely to use the SPK decision model than perform probability matching (74.2%, *P*\* > 0.999), were most likely to use both sensory and reward information to guide their decisions (63.8%, *P*\* > 0.999), and were likely to have lapses (73.0%, *P*\* = 0.998).

**Figure 9:**
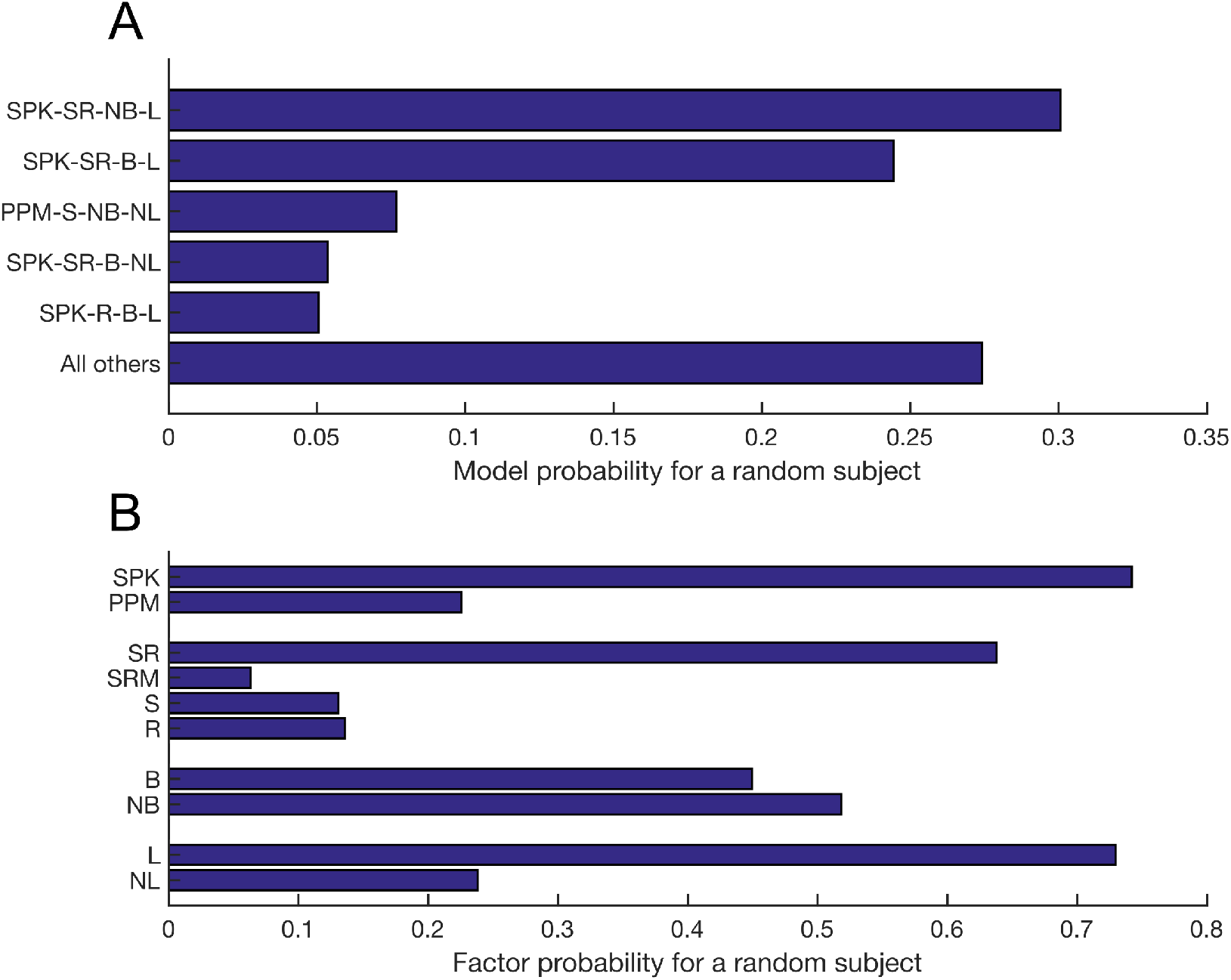
Model selection results for Experiment 2. Same conventions as Fig. 5. (A) The two top models, ‘SPK-SR-NB-L’ and ‘SPK-SR-B-L’, are identical in three of four factors and only differ in whether or not they include decision bias. The third-best model, ‘PPM-S-NB-NL’, only has one free parameter (*α_s_*) and therefore is minimally penalized in the calculation of *BIC* from *NLL*. (B) There was significantly more support for SPK models (‘SPK’ vs. ‘PPM’; *P*\* > 0.999), models using both sensory and reward information (‘SR’ vs. others; *P*\* > 0.999), and models with lapses (‘L’ vs. ‘NL’; *P*\* = 0.998). Models with and without decision biases were approximately equally likely in the population.

Given that decision bias was the only factor not to reach population-level significance and that the top two models were identical other than the presence or absence of decision bias, we further examined the extent to which subjects’ biases influenced model fits. First, we collapsed our family of 32 models across the bias dimension to create a group of 16 model-sets, where each model-set contained both of the models that were identical in the other three factors (e.g., ‘PPM-SRM-B-L’ and ‘PPM-SRM-NB-L’ were combined into the model-set ‘PPM-SRM-?-L’). BMS applied to these 16 model-sets clearly indicated that the set ‘SPK-SR-?-L’ was significantly more likely to be found in the population (54.5%, *P*\* > 0.999). Next, we looked to see if there was a relationship between the bias parameter ***π*** from the full model ‘SPK-SR-B-L’ and the change in model evidence Δ*BIC*_bias_ between the top two population-level models. Positive Δ*BIC*_bias_ means that evidence is stronger for the model without bias, ‘SPK-SR-NB-L’. We used the entropy of ***π*** as a measure of decision bias for each subject:

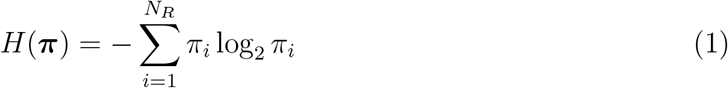

For any discrete distribution, the uniform distribution maximizes entropy (for *N_R_* = 3, ***π*** = [1/3 1/3 1/3], and *H*(***π***) = 1.585 bits), and any deviations from this distribution will decrease entropy. Furthermore, if ***π*** is the discrete uniform distribution, then inclusion of the decision bias parameter in the model will result in no change to the model *NLL* but will increase the *BIC* due to the addition of extra free parameters. Therefore, we hypothesized that decreasing entropy (i.e., increasing decision bias) would be associated with lower *BIC* (i.e., better performance) on the model with decision bias relative to the model without decision bias.

We found a strong correlation in the population between a subject’s entropy and their Δ*BIC*_bias_ (Fig. 10; *r* = 0.668, *p* < 0.001). However, there appeared to be outliers in the population distribution of entropies, and the distribution of entropies was best fit as a mixture of three Gaussian distributions (*BIC*_3_ = −20.83, compared to *BIC*_2_ = −18.49 and *BIC*_4_ = −14.47). Therefore, to ensure that this correlation was not driven by outliers in the data, we also ran correlations over different subsets of subjects. Specifically, in our main cluster of *n* =18 subjects (black stars in Fig. 10), we still found a strong correlation between entropy and Δ*BIC*_bias_ (Fig. 10, inset; *r* = 0.632, *p* < 0.005). This correlation persisted even when only two clusters were used to partition subjects (black stars and blue squares combined, *n* = 22 subjects; *r* = 0.515, *p* = 0.014).

**Figure 10:**
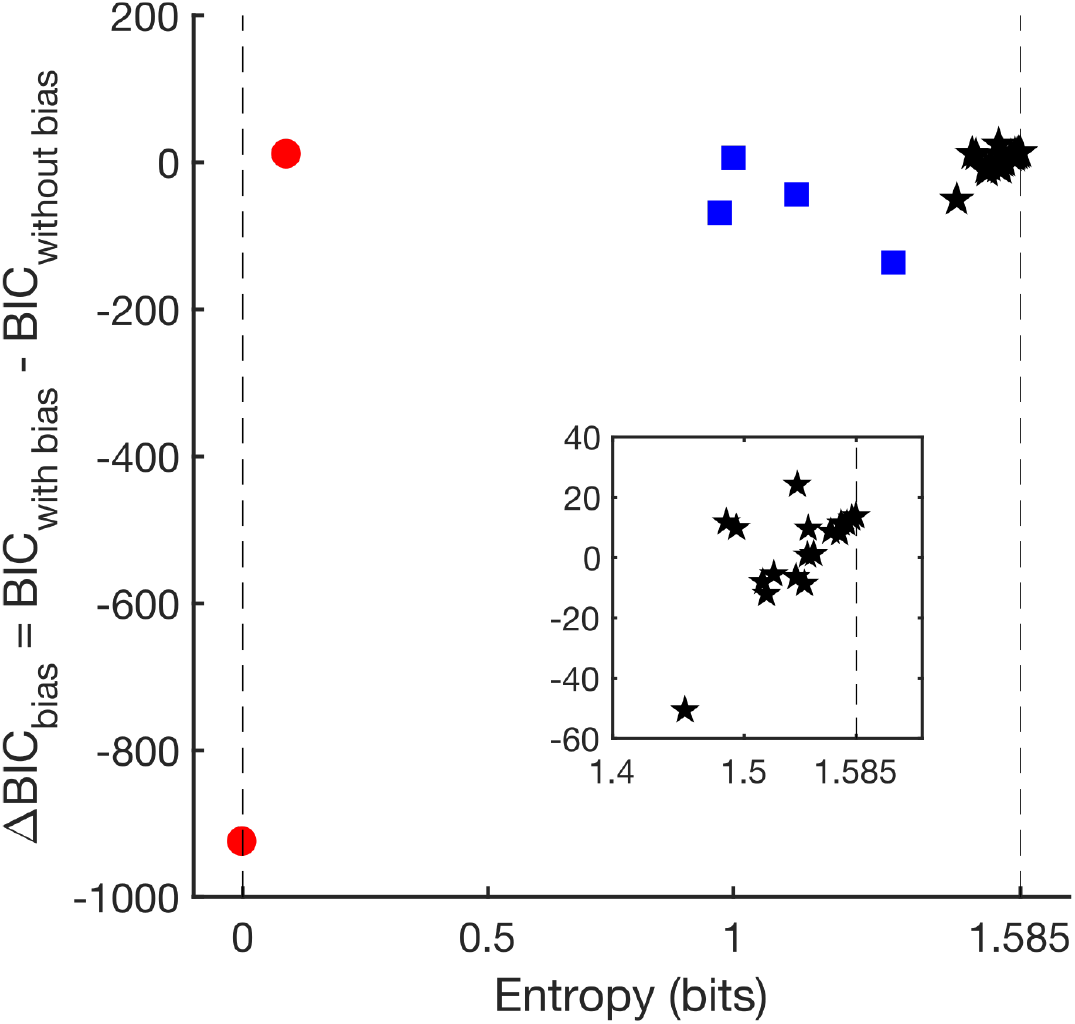
Correlation between decision bias and model fits. Across the population, the relative evidence for ‘SPK-SR-NB-L ‘ vs. ‘SPK-SR-B-L’ was correlated with the entropy of decision bias parameter in the ‘SPK-SR-B-L’ model, 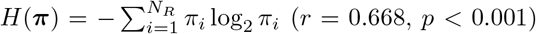. Δ*BIC* > 0 means evidence is stronger for ‘SPK-SR-NB-L’, and Δ*BIC* < 0 means evidence is stronger for ‘SPK-SR-B-L’. Entropy decreases as decision bias increases, is minimized when subjects only ever pick one rule (dashed line on left), and is maximized when ***π*** is the discrete uniform distribution (dashed line on right). The distribution of entropies was best fit by a Gaussian mixture model with three components (black stars, blue squares, red circles), and the correlation was still found when only applied to the main cluster (inset, black stars; *r* = 0.632, *p* < 0.005; see main text for details).

All 32 behavioral models in the family (and the ‘SensoryChaser’ control model) predicted rule selections significantly better than the ‘BiasOnly’ model (Fig. 11; one-tailed t-test, Bonferroni-Holm correction for multiple comparisons), suggesting that subjects were using incoming sensory and/or reward information to guide their decisions. As in Experiment 1, the model with maximal complexity, ‘SPK-SR-B-L’, was the best at predicting rule selections across subjects (27.07± 15.9% above chance). Mean prediction improvements for each model were correlated with the number of parameters in the model (*r* = 0.674, *p* < 0.0001) but not the BMS posterior probability for the model (*r* = 0.218, *p* = 0.22). As expected from the BMS factor results, SPK models predicted rule selections better than PPM models (paired *t*(15) = 4.76, *p* < 0.001) and models with lapses predicted better than models without lapses (paired *t*(15) = 4.04, *p* < 0.005). Additionally, models with decision bias predicted better than those without (paired *t*(15) = 15.3, *p* < 0.0001), and models using both sensory and reward information predicted better than models using only one information source (one-way ANOVA on information source, *F*(3,28) = 17.9; post-hoc Tukey’s test on pairwise combination of ‘SR’ or ‘SRM’ vs. ‘S’ or ‘R’, all *p* < 0.001). In the case of every pattern, as expected, more complex models were better at predicting behavior than less complex models.

**Figure 11:**
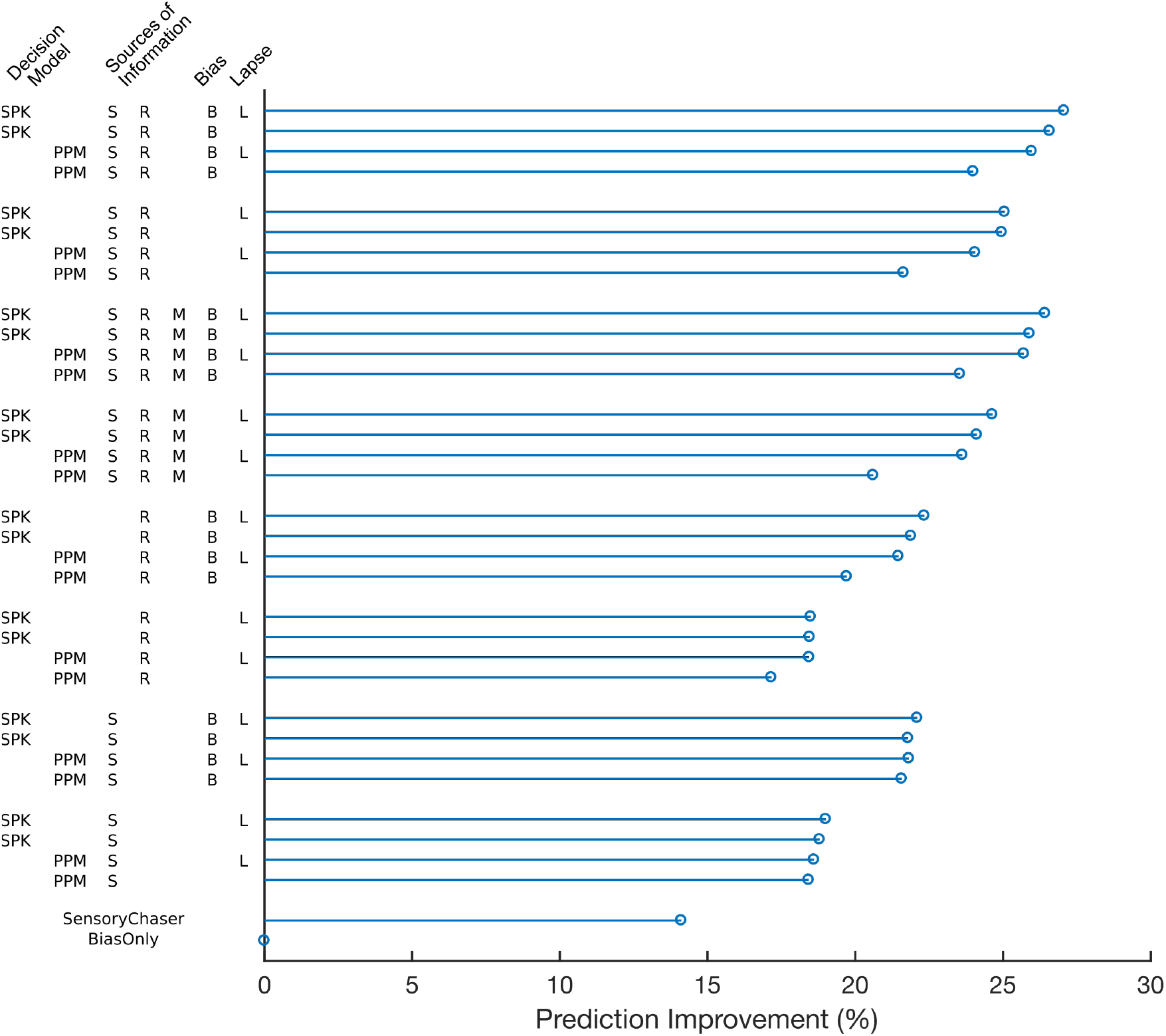
Prediction improvement for Experiment 2. Average prediction improvement was significantly above chance for all 32 behavioral models, and was correlated with model complexity (*r* = 0.674, *p* < 0.0001) but not BMS posterior probability (*r* = 0.218, *p* = 0.22).

We also compared prediction improvements to see if rule selections in Experiment 2 were more or less predictable than those from Experiment 1. We found that, across our population of models, models on average predicted rule selections better when applied to Experiment 2 data than when applied to Experiment 1 data (Fig. 12; paired *t*(31) = 6.13, *p* < 0.0001). Only five models individually fit Experiment 1 data better on average: ‘PPM-R-NB-NL’, ‘PPM-R-NB-L’, ‘SPK-R-NB-NL’, ‘SPK-R-NB-L’, and ‘PPM-R-B-NL’. Of these five models, all of them only use reward information (‘R’) and 4/5 do not have decision bias (‘NB’). Using reward information only (compared to using both sensory and reward information) was associated with inferior prediction in Experiment 2 but not in Experiment 1.

**Figure 12:**
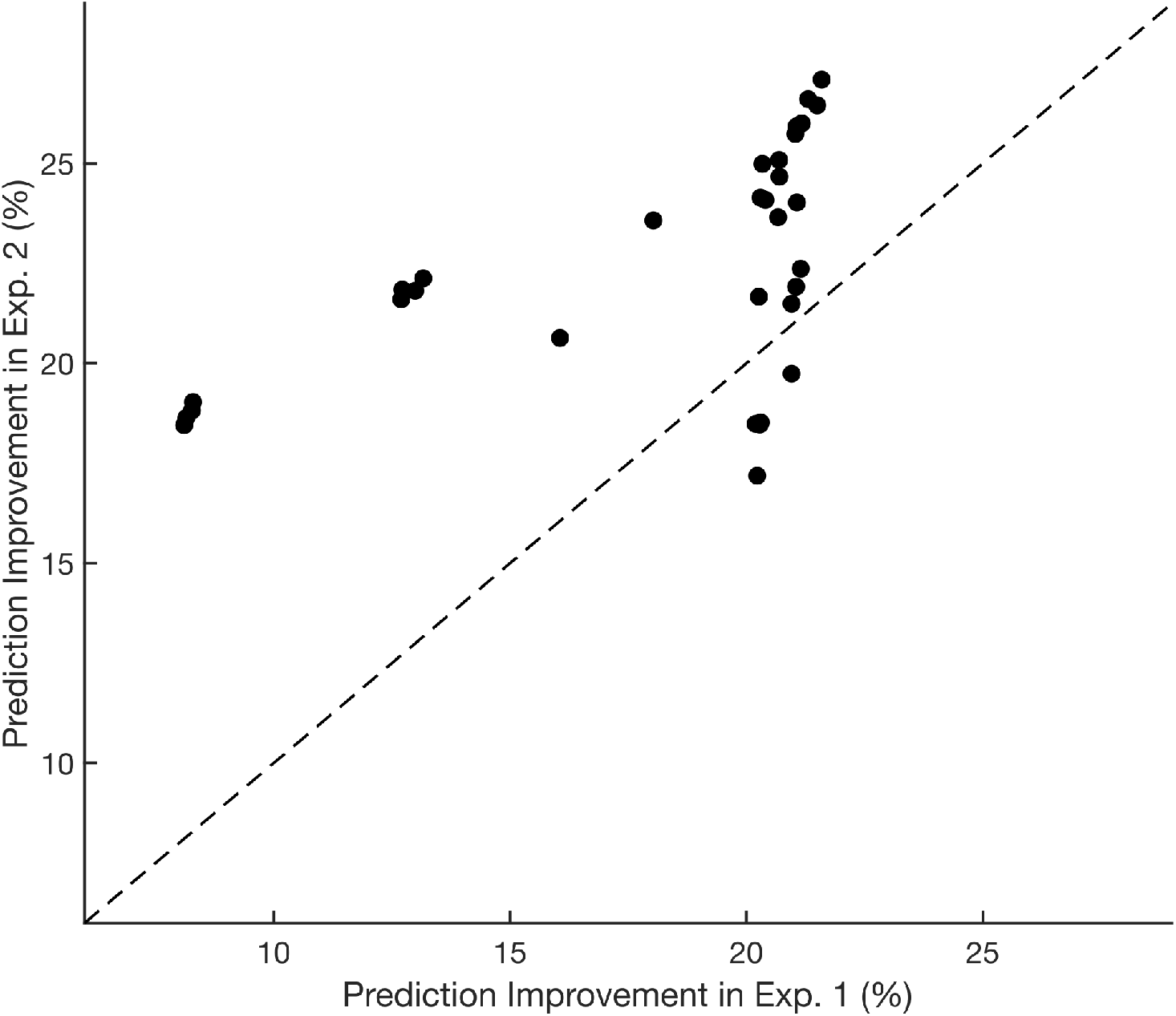
Models were more successful at predicting rule selections in Experiment 2 than in Experiment 1 (paired *t*(31) = 6.13, *p* < 0.0001), represented by data points lying above the dashed unity line. Each data point represents prediction improvement for a single model, averaged across all subjects within an experiment. Only 5/32 models predicted rule selections better on Experiment 1.

Overall, the results of Experiment 2 suggest that subjects utilized both sensory and reward information to guide their rule selection. Interestingly, models that only incorporated sensory history and models that only incorporated reward history were approximately equally poor at explaining the behavior of subjects. Taken together, this indicates that, in tasks where sensory is more useful, sensory information is more likely to be used in conjunction with reward information, and both model-based and model-free learning may be used in concert to generate behavior.

While no individual model was significantly better than all others, the model-set ‘SPK-SR-?-L’ did outperform all other model-sets. At the level of factors, models with stochastic posteriors outperformed models that implemented probability matching, suggesting that subjects had very little noise added to their decision variable. Other groups have also recently found evidence for stochastic posterior models [9], particularly when task performance is based on fine motor behaviors (e.g., using a joystick as opposed to pushing a button) and sensory cues of reward probability are present and accurate [1].

We did not find a consistent effect of decision bias in our population. However, we did find that subject differences in evidence between the top-performing models with and without bias were strongly related to subject biases. Additionally, most of our subjects had small levels of bias, although there did appear to be two distinct groups of outliers with larger biases. One potential explanation for this is that some subjects were not properly motivated to maximize their performance for some or all of the behavioral session, and therefore developed rule preferences. Even with our small-bias group, though, we found predictable heterogeneity in model fits as a function of bias. Given the relatively large number of trials performed by each subject (≈ 1000), we believe that these empirical biases are not simply artifacts of taking a finite number of samples from an unbiased (i.e., uniform) distribution. Therefore, we suggest that the benefits of including bias parameters in the model outweigh the risks of overfitting rule choices.

## 3. Discussion

The behavioral tasks used in Experiments 1 & 2 shared a fundamental structure (preselect a rule and then use it to flexibly behave) but differed in the specific rules, stimuli, and motor behaviors used. These lower-level task changes led to drastic changes in the behavioral strategies used by human subjects to make their rule selections. In Experiment 1, subjects had to choose between two rules that existed in different perceptual dimensions and report a discrimination using button presses: subject behavioral strategies were heterogeneous but typically dependent on reward history. In Experiment 2, subjects had to choose between three rules that existed in the same perceptual dimension and use a joystick to aim for the desired target: subject behavioral strategies were more homogeneous and dependent on both sensory and reward history.

### 3.1. Estimating task performance

One possible explanation for the difference between the findings is that it was easier to estimate task performance from sensory stimulus features in Experiment 2 than in Experiment 1. Expected performance on any task or rule with binary outcomes (where *O* =1 represents correct trials and *O* = 0 represents errors) is:

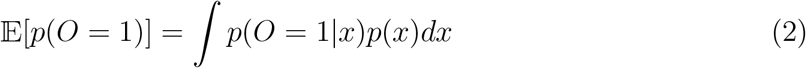

where *p*(*O* = 1|*x*) is the psychometric curve (i.e., the relationship between stimulus features and reward probabilities), *p*(*x*) is the generative distribution for stimulus features, and all probabilities are conditioned on the task or rule (which is dropped from the notation for the sake of clarity). Therefore, comparing the expected difficulties of multiple tasks requires the estimation of both the psychometric curve and generative distribution for each candidate task. If, however, psychometric curves for the tasks are the same, then only one psychometric curve needs to estimated. In Experiment 1, psychometric curves for brightness discriminations and size discriminations are distinct and therefore both must be estimated separately in order to estimate task difficulty without using heuristics. When confronted with these additional task demands, our results indicate that humans instead default to heuristics that are reliant only on trial outcomes and not on the generative distribution *p*(*x*). In contrast, in Experiment 2, the probability of hitting a target as a function of target size should be independent of target color, so only one psychometric curve needs to be estimated. As a result, subjects in Experiment 2 used both sensory and reward history to guide their rule selections.

We found that models without lapse parameters best explained subject behavior in Experiment 1, whereas models with lapse parameters performed best in Experiment 2. We believe that this effect is driven by the number of rules used in each task. A subject with both decision bias and lapses can be approximated by a model with only decision bias, and the quality of this approximation degrades as the lapse rate increases and as the number of possible rules increases. Therefore, the increase in model fit quality due to explicitly modeling lapses is greater in Experiment 2 (*N_R_* = 3) than in Experiment 1 (*N_R_* = 2), which compensates for the *BIC* penalty associated with adding a model parameter. Moreover, when subjects are faced with more rule options, they may be more likely to forget which rule they were planning to choose, leading to more instances of random selection.

### 3.2. Temporal dependence and probability matching

We also found that stochastic decision-making models outperformed probability matching models in Experiment 2, but found no evidence in favor of either class of decision model in Experiment 1. One possible explanation of this effect is based on motor noise and temporal independence. Noise in the motor system is well-modeled as being independent of past motor behavior [27], and when trial outcomes are based on motor behaviors (or appear to be based on them), this temporal independence leads to more optimal choices instead of suboptimal probability matching [1]. In particular, Green et al. (2010) postulated that subjects typically assume that outcomes in nature are temporally dependent (e.g., the tree that had fruit yesterday is likely to have fruit today), and showed that a mismatch between this expectation of temporal dependence and the actual temporal *independence* that is typically built into experimental tasks drives probability matching behavior. However, when subjects are given the illusion that their motor behavior affects trial outcomes (even when it does not actually), this expected source of temporal independence allows them to account for the temporal independence that is built into the experimental task, and more optimal behavior emerges [1]. In line with these findings, subjects in Experiment 2, which required fine motor behaviors that are sensitive to temporally independent noise (moving a joystick in a precise direction), may have been more likely to assume errors were due to motor noise and not a misunderstanding of the relationship between stimulus features and task difficulty. This would lead to less exploratory behavior (such as probability matching) and therefore more optimal decisions. In Experiment 1, where subjects reported decisions with gross motor actions that are robust to noise (pushing a button), subject behavior was more heterogeneous with no population-level preference for either stochastic posterior or probability matching models, suggesting that some subjects believed they understood the relationship between stimulus features and task difficulty, while others were less confident.

### 3.3. Overall interpretations and discussion

Here, we turned to computational modeling in an attempt to better understand serial decision-making. Specifically, we utilized the factorial approach [9, 24] in combination with Bayesian model selection [25] to test a large number of behavioral models. We believe that this approach is superior to traditional behavioral modeling techniques where a single model framework is selected, and then the parameters of that model are fit to subject data. This approach, for example, is able to show that many of our human subjects were best fit by a number of different behavioral models (Figs. 4 and 8), suggesting that humans use a variety of strategies to solve the same task. If we had pre-selected only one behavioral model for data fitting, we would have missed out on this insight.

Moreover, computational models of behavior are elegant ways to summarize many hundreds of trials worth of data. For example, our most complex behavioral model features seven parameters, effectively allowing us to summarize the data from each subject as one point in 7-dimensional space. By applying the same behavioral model to data from subjects performing further variations of our serial decision-making task, we can see how subtle changes to the task lead to changes in the behavioral strategies used. If the same subjects perform multiple tasks in a behavioral session, then we can use model comparison to investigate how a subject’s behavioral strategy on one task relates to behavior on other tasks, by examining both the dominant strategy for that subject (as defined by *BIC*) and the parameter fits for that model. Computational models, by trying to learn the strategies that generate behavior, allow for a more fine-grained comparison than simply comparing bulk metrics such as performance.

In summary, we used normative probabilistic models to determine the behavioral strategies used by humans to select between uncertain options. When sensory features span different dimensions, humans resort to a model-free learning strategy that relies exclusively on reward history. However, when sensory features can be compared more intuitively, humans adopt a hybrid strategy that relies on both reward-driven model-free learning and sensory-driven model-based learning. Furthermore, when trial outcomes are determined by subjects beliefs regarding motor noise (in addition to sensory noise), subjects exhibit reduced decision noise and make more optimal decisions.

## 4. Methods

### 4.1. Behavioral models

Our objective was to develop a model that is motivated by the optimal behavior of an ideal observer and then investigate ways of sensibly perturbing the model in order to capture suboptimalities in real-world behavior and strategy. Our core behavioral model featured three modules: a Sensory Module, which tracked incoming sensory information; a Reward Module, which tracked reward histories for each rule; and a Decision Module, which integrated information from the Sensory Module and Reward Module with potential biases and lapses.

#### 4.1.1. Sensory Module

The purpose of the Sensory Module was to recursively track the distributions of sensory features (the distance from which a basketball or soccer shot must be taken, the size/brightness/color of an object, etc.) for each rule *i* on each trial *t* according to the following update rules:

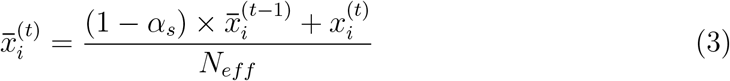

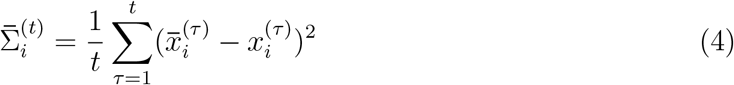

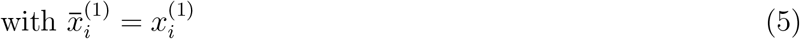

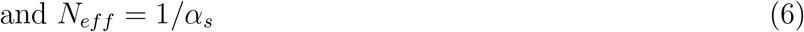

where 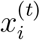 is the observed feature for rule *i* on trial *t*, 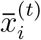 and 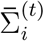 are the mean and variance of the feature distribution for rule *i* incorporating all evidence through trial *t*, and the free parameter *α_s_* is the sensory learning rate. The learning rate parameter was included to allow the mean to vary dynamically rather than forcing it to converge to a stable estimate. In contrast, the estimate of the variance around that dynamically changing mean was assumed to be constant and to converge in the limit of infinite observations. Then, the predicted feature for rule *i* on trial *t* + 1 was drawn from the predictive distribution:

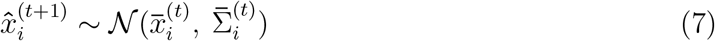

All features (size and brightness differences in Experiment 1, target size in Experiment 2) were such that greater feature strengths are monotonically associated with greater task performance (e.g., as the difference in brightness between two targets increases, it becomes easier to discriminate which target is less bright). Therefore, an optimal decision-maker would be interested in the probability that a random draw from the feature distribution for rule *i* is greater than random draws from feature distributions for all other rules *j* ≠ *i*. Utilizing the closed-form solution for the difference between normally-distributed random variables, dropping the trial superscript for clarity, and setting the numbers of possible rules to 2, yielded:

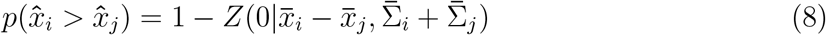

where *Z*(·) is the cumulative distribution function of the normal distribution.

A rational decision-maker should base their choices systematically on the relative likelihoods that a target of a particular color will be largest. Therefore, we employed the power function approximation to the stochastic posterior model (SPK; 9) to model decision-making. The SPK model is of the form:

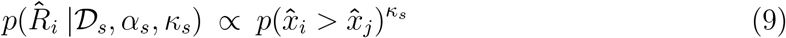

where 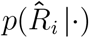 is the model probability of picking rule *i* given data and parameters, 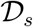 is the sensory stimulus history, and the power exponent *κ_s_* is a free parameter controlling how noisy or variable the decision model is. One primary benefit to the SPK model is its versatility; by changing the value of *κ_s_*, the model can implement probability matching (*κ_s_* = 1; 9) or maximum exploitation (*κ_s_* → ∞), and can even ignore predictive feature distributions (*κ_s_* = 0) or preferentially select harder rules (*κ_s_* < 0).

#### 4.1.2. Reward Module

Analogous to the Sensory Module, the purpose of the Reward Module was to recursively track distributions of rewards for each rule *i* on each trial *t* according to the following update rules:

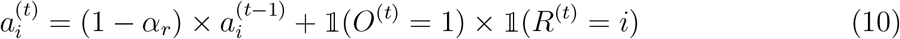

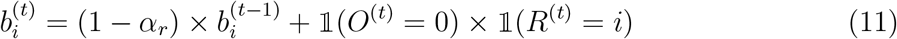

where 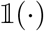 is the indicator function, *O*^(*t*)^ is the actual outcome of trial *t* (1 for correct, 0 for error), *R*^(*t*)^ is the actual rule on trial *t*, and the free parameter *α_r_* is the reward learning rate. Then, the predicted reward likelihood *Ô* for rule *i* on trial *t* + 1 was drawn from the predictive distribution:

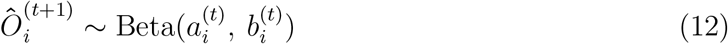

Also analogous to the Sensory Module, we were interested in determining the probability that a random draw from the reward distribution for rule *i* is greater than random draws from reward distributions for all other rules *j* ≠ *i*. However, the difference between beta-distributed random variables does not have a closed-form solution. Therefore, we substituted in the geometric mean of the beta distribution *Õ_i_*, again dropping the trial superscript for clarity and utilizing the SPK model:

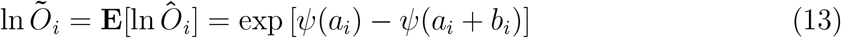

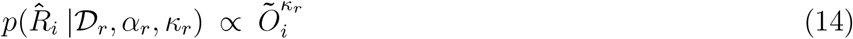

where *ψ* is the digamma function, 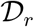 is the reward history, and the power exponent *κ_r_* is the free SPK parameter.

#### 4.1.3. Decision Module

The Decision Module took the outputs of the Sensory Module and Reward Module and combined them with biases and lapses to ultimately determine the probability that the model selected rule *i* on any given trial:

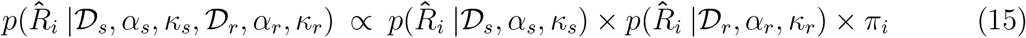

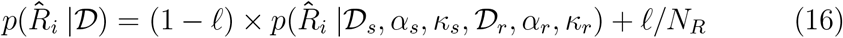

where the free vector ***π*** contains the rule biases, 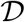 is a shorthand representation for all data and parameters supplied to the model, the free parameter *ℓ* is the lapse rate, and *N_R_* is the number of rules in the task. The final output of the model was 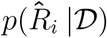, the probability of the model selecting rule *i* on any given trial.

#### 4.1.4. Creating a family of models

The final model had 5 + (*N_R_* − 1) free parameters: *α_s_, κ_s_, α_r_, κ_r_, ℓ*, and the bias vector ***π***, which has *N_R_* − 1 free parameters because it is subject to the constraint 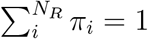. We employed a factorial approach to generate a large family of models [9, 24] by setting different subsets of free parameters to be equal to constants. Our family of models was therefore the combinatorial set of the following factors and their levels:

1. *Decision Model* (2 levels): stochastic posterior model (‘SPK’: *κ_s_* and *κ_r_* are free) or posterior probability matching (‘PPM’: *κ_s_* = 1 and *κ_r_* = 1)
2. *Sources of Information* (4 levels): both sensory and reward information (‘SR’: *α_s_*, *κ_s_*, *α_r_*, and *κ_r_* are all free), sensory information only (‘S’: *α_r_* = 0 and *κ_r_* = 0), reward information only (‘R’: *α_s_* = 0 and *κ_s_* = 0), or both sources with matched learning rates (‘SRM’: *α* = *α_s_* = *α_r_*)
3. *Decision Bias* (2 levels): present (‘B’: ***π*** is free) or absent (‘NB’: all *π_i_* = 1/*N_R_*)
4. *Lapse* (2 levels): present (‘L’: *ℓ* is free) or absent (‘NL’: *ℓ* = 0)

Therefore, each of the 2 × 4 × 2 × 2 = 32 models in this family can be uniquely described by a string. For example, the full model described above is ‘SPK-SR-B-L’ and, for *N_R_* = 2, has 6 free parameters. In contrast, a model that used the stochastic posterior model, only utilized sensory information, had decision bias, and had no lapse would be denoted as ‘SPK-S-B-NL’ and, for *N_R_* = 2, has 3 free parameters (*α_r_* = 0, *κ_r_* = 0, and *ℓ* = 0).

We also implemented two control models: ‘BiasOnly’ and ‘SensoryChaser’. The ‘BiasOnly’ model has *N_R_* − 1 free parameters, the rule bias vector ***π***, and used the static decision model:

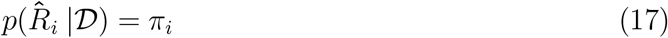

on every trial. The ‘SensoryChaser’ model has one free parameter, the lapse parameter *ℓ*, and simulates the behavior of a subject who selects the rule that had the greatest feature strength (i.e., the easiest rule) on the immediately previous trial with probability 1 − *ℓ* and doesn’t utilize any other information. This corresponded to the dynamic decision model:

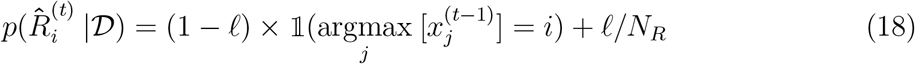

#### 4.1.5. Fitting and evaluating models

We evaluated model performance using three primary metrics, the negative log-likelihood (*NLL*), the Bayesian information criterion (*BIC*), and prediction accuracy (how often the model selects the rule that the subject actually selected):

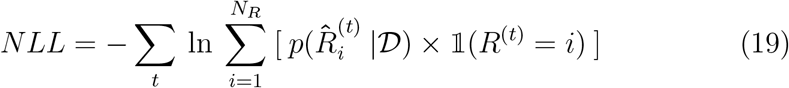

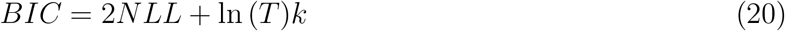

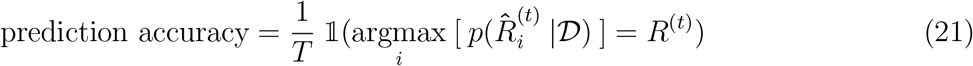

where *T* is the number of trials the subject performed and *k* is the number of free parameters in the model. We used *BIC* rather than *AIC* to decrease the likelihood of selecting overparametrized, more general models, which can be a particular problem with a family of nested models [28].

All models were fit eight times per subject with randomized initial conditions in an attempt to avoid local minima. Models were fit using Matlab’s `fmincon` function with a tolerance of 10^−5^, and the individual run that yielded the lowest *NLL* was retained for further analysis. Because all models in the family were formulated by starting with the ‘SPK-SR-B-L’ model and setting free parameters as constants, and adding parameters to a model (i.e., increasing model complexity) will never result in a worse fit to data (i.e., increasing the NLL), we determined that the ‘SPK-SR-B-L’ model should yield the lowest *NLL* for every subject. Therefore, we utilized the *BIC* to facilitate comparison between models of varying complexity. For any given subject, the number of trials performed *T* is constant, so adding one additional parameter to the model increases the *BIC* by ln (*T*). Lastly, we were interested in prediction accuracy as a measure of how well the models can predict the actual rule selections made by the subject. Specifically, we are interested in the prediction improvement for each model over chance, where chance is prediction accuracy for the ‘BiasOnly’ model.

We also used Bayesian model selection (BMS) to directly compare between models [25]. BMS computes the evidence for each model based on marginal likelihoods from each subject (or an approximation to the marginal likelihood, such as (−1/2)*BIC*). Specifically, the BMS algorithm is a variational Bayesian method that treats different models as random variables in a Dirichlet distribution while allowing for in-group heterogeneity (i.e., treating subject identity as a random effect). BMS provides three key related metrics that allow formal comparison between models: the parameters of the fit Dirichlet distribution, the multinomial distribution over models that defines the likelihood that each model generated the data of a random chosen subject, and the “exceedance probability” *P*\* that one model (or one factor level) is more likely than any other. We initialized the BMS algorithm with a symmetric Dirichlet distribution and *α*_0_ = 0.25, which corresponds to the weak belief that only a few behavioral models are actually present in the population [9].

### 4.2. Experiment 1

Twenty-four naïve human subjects, with normal or corrected-to normal vision, were recruited from the Duke University community. All individuals were older than 18 years and gave informed consent through protocols approved by the Duke Institutional Review Board. Subjects in Experiment 1 completed an average of 465 ± 70.5 trials in an hour-long behavioral session.

#### 4.2.1. Behavioral task

Behavioral sessions consisted of blocks of 30-60 trials. Transitions between blocks were continuous, covert, and uncued. Within a single block, rule difficulties were globally coupled: one rule was designated as the “easy” rule and the other rule was necessarily designated as the “hard” rule (these assignments swapped at the start of every new block). Importantly, the difficulty distributions were overlapping. This meant that the rule difficulties were locally uncoupled, such that a single trial could, for example, feature both a difficult size discrimination and a difficult brightness discrimination.

At the start of each trial, the size (measured in pixels) and brightness (measured in screen brightness units ∈ [0, 1]) of the target on the left were drawn from normal distributions:

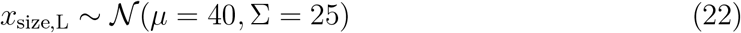

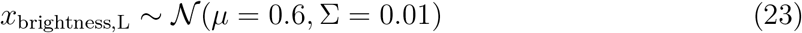

The size and brightness of the target on the right was defined relative to the target on the left. Specifically, “easy” and “hard” distributions were defined by the ratio of sizes or brightnesses between target on the right and the target on the left. Both difficulty distributions were mixtures of the same two discrete uniform distributions, one with possible R:L ratios of 0.7:1, 0.8:1, 1.2:1, or 1.3:1 (corresponding to easy discriminations), and a second with possible R:L ratios of 0.96:1, 0.98:1, 1:1, 1.02:1, or 1.04:1 (corresponding to difficult discriminations). The “easy” rule had R:L ratios drawn predominantly from the easier distribution (*π*_easy_ = 0.9, *π*_dfficuit_ = 0.1). The “hard” rule had R:L ratios drawn predominantly from the more difficult distribution (*π*_easy_ = 0.1, *π*_difficult_ = 0.9)

As stated above, these size differences and brightness differences are measured in different units. As a result, some sort of scaling factor must be used to convert between size differences and brightness differences before they can be directly compared in Equation 8. To determine the optimal scaling factor, we used Matlab’s `fminsearch` function to minimize the divergence (as a function of the scaling factor) between two distributions: the raw distribution of size differences and the distribution of scaled brightness differences. We used the symmetrised KL divergence as our divergence metric to be minimized:

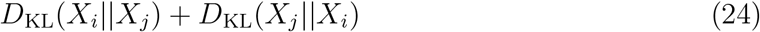

This procedure yielded a scaling factor of 70.3, and *x*_brightness_ values were multiplied by this factor before being input into Equation 3.

### 4.3. Experiment 2

Twenty-four naïve human subjects, with normal or corrected-to normal vision, were recruited from the Duke University community. All individuals were older than 18 years and gave informed consent through protocols approved by the Duke Institutional Review Board. Subjects in Experiment 2 completed an average of 952 ± 164.2 trials in an hour-long behavioral session.

#### 4.3.1. Behavioral task

Like the task used in Experiment 1, the task used in Experiment 2 required subjects to first choose a behavioral rule, and then use that rule to flexibly guide behavior in response to complex stimuli. Specifically, we had subjects perform a “shooting” task that required them to use a joystick to pre-select a color, and then shoot at a target of that color among multiple distractors (Fig. 7). A video of the task can be found at https://goo.gl/iXZybV.

In the rule-selection stage (Fig. 7A, left), subjects were shown a ring that was equally split into three distinct colored “rule” regions (red, green, and blue). Subjects were free to select one of the colored regions at any time after their onset by directing a joystick radially in the direction of their chosen target. The ring target was always centered on the screen, but the precise boundaries between the three rule regions varied randomly from trial-to-trial, such that the location of one particular rule region was unpredictable on any given trial. After the joystick returned to its resting position, the trial automatically progressed to the rule-implementation stage.

In the rule-implementation stage (Fig. 7A, right), subjects were shown a small white ball (10px diameter) at the center of the screen, and three colored “arc” targets (red, green, and blue) at the same radial distance away from the ball (set to 0.3 times the height of the screen in pixels, for a landscape-oriented monitor). The targets varied in their arc-widths and their locations, but their centers were always equally spaced around the circumference of a circle (120° apart). Importantly, the location of each target was random from trial-to-trial and could not be predicted by the location of the colored rule region in the rule-selection stage. The subjects’ goal was to use the joystick to “shoot” the ball at the target that was the same color. To determine the ball’s trajectory, we determined when the subject initially began to displace the joystick from its resting position, waited 50 ms, and then sampled the new joystick position. The resting joystick position was subtracted off from the sampled joystick position, the difference was converted into an angular trajectory, and the ball began moving according to that trajectory. In practice, this trajectory was typically modified by some correction factor before visual feedback began (see *Controlling motor error* below for details). If the ball’s angular trajectory was within the angular bounds of the target, the ball collided with the target, stopped moving, and an auditory cue sounded. If the they did not successfully hit the target (or hit an incorrect target), the ball continued moving until it hit the edge of the screen and no auditory cue sounded.

As in Experiment 1, behavioral sessions consisted of blocks, and transitions between blocks were continuous, covert, and uncued. Within a block, one colored rule was designated as the “easy” rule (i.e., largest target size, on average), one was designated as the “medium” rule, and the third was designated as the “hard” rule (i.e., smallest target size, on average; Fig. 7B). Blocks lasted 6-9 trials, and at the end of each block, there was a 40% chance of swapping the identities of the “easy” and “medium” rules, a 40% chance of swapping the identities of the “medium” and “hard” rules, and a 20% chance of the rankings staying the same (Fig. 7C). A single rule could never transition directly from “easy” to “hard” or vice versa in a single block transition. On a trial-by-trial basis, target sizes were drawn from the following distributions:

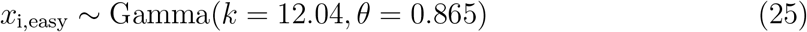

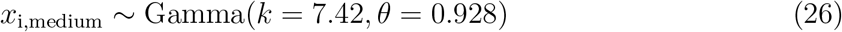

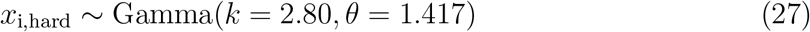

This yielded target size distributions with means of 10.41/6.88/3.97° and standard deviations of 3.00/2.53/2.37° (E/M/H, respectively). These parameters were selected to yield performances (i.e., target hit rates) of 0.6/0.425/0.25 (E/M/H, respectively) while maximizing overlap between the distributions, given an ideal subject motor error of *σ*_ideal_ = 6px.

##### Controlling motor error

At the start of every session, subjects completed 80 “calibration” trials. These trials mimicked the rule-implementation stage of the main task, but with only a single 7.5° wide white target in the periphery instead of three colored targets of variable size. The purpose of these calibration trials was to generate an initial estimate of each subject’s motor error, *σ*_actual_. Then, during the actual testing phase, we attempted to equalize performance across subjects through the following procedure:

1. We calculated the correction factor *f* = *σ*_ideal_/*σ*_actual_.
2. On each trial, we determined the angular deviation *d* between the actual trajectory and the center of the chosen target.
3. If |*d*| < 45°, we shifted the trajectory such that the new shifted deviation *d*′ = *f* × *d*. if |*d*| ≥ 45°, no shift was applied and *d*′ = *d*.
4. The deviation d’ was used for both visual feedback (the trajectory shown on screen) and reward feedback (whether or not the ball hit the target).
5. Every 50 trials (after the initial 100 trials had elapsed), the motor error *σ*_actual_ was updated to control for practice effects and motor learning.

In practice, actual motor errors were always greater than *σ*_ideal_ = 6°, meaning that the shifting procedure increased subject accuracy. All subjects who commented on this increase in accuracy attributed it to the successful “calibration” of the joystick, rather than any experimental manipulation of trajectories. Additionally, actual deviation magnitudes were almost always < 45° unless subjects purposefully aimed for a different target. No subjects commented on any perceived misalignment between intended and displayed target trajectories.

## 5. Acknowledgments

This material is based upon work supported by the National Science Foundation under Grant 1539687 to M.A.S. The authors declare no competing financial interests.

